# Growth-promoting rhizobacteria amend the defense of strawberry plants against sequentially attacking herbivores

**DOI:** 10.1101/2025.11.16.688721

**Authors:** Afsane Hosseini, Mojtaba Hosseini, Peter Schausberger

## Abstract

Plant defense systems such as induced resistance (IR; induced by herbivores) and induced systemic resistance (ISR; induced by beneficial rhizobacteria) are modulated by the same signaling pathways within plants and hence interact. ISR allows enhanced protection of aboveground plant parts against herbivores before IR induction. Both ISR and IR are systemic and may involve the production of toxic, antifeedant and/or repellent compounds, and/or reduce nutrient availability, which may in consequence affect plant usability and palatability for later arriving herbivores. The combined effects of ISR and IR on different herbivores sharing the same plant and plant performance have been rarely addressed. Here, we assessed the effects of three plant-growth-promoting rhizobacteria (PGPR), *Azotobacter chroococcum*, *Azospirillum brasilense* and *Pseudomonas brassicacearum,* on the defense response and physiology of strawberry plants upon sequential attack by two herbivores with different feeding modes, two-spotted spider mites *Tetranychus urticae* and cotton aphids *Aphis gossypii*. Attack of strawberry plants by spider mites and aphids adversely affected the abundance of the later arriving herbivore, mediated by the host plant’s defense system. First-attacking spider mites exerted much stronger adverse effects on later attacking aphids than first-attacking aphids on later attacking spider mites. In absence of PGPR inoculation, the herbivores, especially first-attacking spider mites, severely impaired host plant physiology and productivity. PGPR inoculation primed the plant’s defense system to attack by spider mites and aphids, allowing the plants to produce more secondary metabolites such as phenols. In consequence, the abundances of both herbivores were lower on PGPR-inoculated plants compared to chemically fertilized and control plants. Overall, our study suggests that PGPR inoculation ameliorates the plant damage caused by sequentially attacking herbivores. Additionally, the PGPRs improve the physiology and productivity, and favorably balance the nutritional state, of strawberry plants.

## Introduction

Herbivory selects for the development of defense strategies in co-evolving plants ^1^. These plant defenses include changes in chemical, physiological and morphological plant traits derived from changes in primary and secondary metabolites. A state of the plant defense system that is elicited upon a challenge from living organisms such as herbivores has been termed induced plant resistance (IR) ^2,3^. Induced resistance is modulated by signaling pathways in which phytohormones play a major regulatory role ^4^. Jasmonic acid (JA) and salicylic acid (SA) are two major phytohormones underlying induced plant defenses triggered by herbivores with different feeding modes ^5,6^. It has been suggested that leaf-chewing herbivores (such as lepidopteran larvae) typically activate JA, phloem-feeding herbivores (such as aphids) typically activate SA, and parenchyma cell-content feeders, such as spider mites, may elicit both signaling pathways though predominantly JA ^7–9^. Herbivore-induced plant response may involve the production of toxic, anti-feedant and/or repellent compounds, and/or reduce nutrient availability, which may in consequence affect plant usability and palatability for later arriving herbivores ^10,11^. Plant metabolic changes in response to herbivory are frequently, but not always, found to cause increased defense against future herbivores ^12,13^. For instance, initial herbivory by two-spotted spider mites *Tetranychus urticae* promotes herbivore-induced plant resistance against sequential infestation by tobacco whiteflies *Bemisia tabaci* ^14^. However, previous herbivore infestation can also facilitate further colonization by later arriving herbivores, which may especially occur when the subsequent herbivores are conspecific ^15–17^. For example, larvae of the western corn rootworm *Diabrotica virgifera* develop better on roots previously attacked by conspecifics ^18^.

Another type of induced plant resistance of aboveground plant parts, dubbed induced systemic resistance (ISR), is mediated by beneficial soil microbes ^4,19,20^. Enhanced protection of aboveground plant parts through ISR can be conferred prior to herbivory, e.g., by application of beneficial soil microbes ^21–23^. This type of induced resistance is considered as plant sensitization and has also been dubbed priming ^4^. Beneficial soil microbes may prime the plant to subsequent attacks by herbivores or pathogens by leading to earlier, faster and/or more intense activation of phytohormones, which may have lower costs for the plant than immediate activation of the defense system upon herbivore attack ^4,24,25,26^. Beneficial soil microbes modulate defense signaling pathways that culminate in the production of secondary metabolites ^22^. The most common type of defensive compounds of secondary metabolites are phenolics, which play major roles in resistance against herbivores and which may accumulate in above-ground plant tissue by below-ground association with beneficial soil microbes ^4,27^.

Among beneficial soil microbes, the widespread plant-growth-promoting-rhizobacteria (PGPR) are well known for their potential as efficacious biological agents against herbivorous plant pests ^22,26^. For instance, rhizobacteria inoculation of strawberry plants decreased the life history performance and population growth of two-spotted spider mites compared to those feeding on N-fertilized plants ^28^. In addition, PGPR may improve plant growth by enhancing nutrient and water uptake ^29^. For example, free-living plant-associated bacteria such as *Azospirillum* spp., *Azotobacter* spp., and/or *Pseudomonas* spp. may promote root and shoot growth of plants by providing N solubles and other nutrients, and the production of phytohormones ^30^, which may benefit plant tolerance but may also enhance the nutritional value of the plant tissue for herbivores ^27^. So, PGPR can indirectly affect herbivore fitness by increased plant nutritional value and/or by triggering ISR ^4,31^.

We previously found that three species of PGPR, *Azotobacter chroococcum* (Ac), *Azospirillum brasilense* (Ab), and *Pseudomonas brassicacearum* (Pb) exert adverse effects on the life history performance and population dynamics of *T. urticae* feeding on strawberry plants ^28^. To achieve a more comprehensive understanding of the consequences of PGPR-induced changes in strawberry resistance against herbivores, here we investigated the effects of these three plant-associated PGPRs on the defense response and physiology of strawberry plants upon attack by two herbivores with different feeding modes, two-spotted spider mites *T. urticae* and cotton aphids *Aphis gossypii*, sequentially arriving at the plant. Spider mites feed on plants by extracting the contents of parenchyma cells, while aphids feed phloem sap. We addressed three major questions concerning the interactions between rhizobacteria, strawberry plants and the two herbivores: (i) how does the interaction between the plant-associated free-living rhizobacteria and the sequence of herbivore arrival on the plant affect the abundance of both herbivores? (ii) do the three PGPRs differ in their effects on the plants and herbivores? (iii) do the rhizobacteria-associated plants have the same or different defense levels after the first and second herbivore attack?

## Materials and methods

### Bacteria preparation

*Azotobacter chroococcum* (Ac), *Azospirillum brasilense* (Ab), and *Pseudomonas brassicacearum* (Pb) used in the experiments were originally obtained from the Plant Pathology Laboratory, Department of Plant Protection, Faculty of Agriculture, Ferdowsi University of Mashhad, Iran ^28^. For routine use, the bacteria were grown on nutrient agar (NA), which consists of peptone (0.5%), beef extract (0.3%) and agar (1.5%). Nutrient broth (NB), which is composed of peptone, yeast extract, vitamin B complex and other nutritional requirements, with 15% glycerol was used for long-term storage at -80°C (Merck KGaA, Darmstadt, Germany).

For the experiment, bacterial cultures were removed from long-term storage and grown on NA until they were ready for use. For each rhizobacterial species, an inoculation loop of bacteria was transferred to NB in 1-litre flasks placed on an orbital shaker with 150 rpm for 48 h at 24 °C. Subsequently, the rhizobacterial suspensions were diluted in distilled water to 2×10^9^ CFUs (colony forming units) per ml. The diluted rhizobacterial suspensions were used to inoculate the strawberry plants ^32^. The strains of the three rhizobacterial species used in this study have previously demonstrated growth-promoting effects on strawberry plants ^28^.

### Strawberry plant culturing and treatments

All plant materials used in this study were acquired and utilized in accordance with relevant institutional, national, and international regulations and guidelines. Seedlings (3- to 4-leaf stage) of strawberry (*Fragaria × ananassa* cv. Selva; Yolaflor Co., Mashhad, IR) were individually grown in 1-litre plastic pots (20 cm high, 17 cm Ø) filled with coconut peat (Kumari Coir Products, Singapore), clay and sand (1:1:1 by volume). Each seedling/pot was covered by a transparent cylindrical fine-mesh cage (70 cm high, 30 cm Ø) and kept in a greenhouse at the Agricultural Faculty, Ferdowsi University of Mashhad at 26 ± 1°C, 65 ± 5% RH and natural daylight. The natural photoperiod was about 14/10 h L/D when the study was conducted (early June to mid-August 2022).

The five plant treatments in the experiment were as follows: inoculation of strawberry roots with (1) *A. chroococcum* (Ac), (2) *P. brassicacearum* (Pb), or (3) *A. brasilense* (Ab), all without chemical fertilizer application. The remaining two treatments were (4) chemical fertilization (without PGPR inoculation), and (5) control (without PGPRs and without chemical fertilizer). Prior to planting, the roots of the strawberry seedlings were soaked for 40 min in aqueous suspensions of either Ac, or Ab, or Pb (density 2×10^9^ CFU/ml) for the PGPR treatments (1), (2), and (3). Subsequently, the seedlings were allowed to air-dry for 30 min, after which they were transplanted into the pots as described before ^28^. The pots were irrigated with tap-water once every 2 d. For the chemical fertilizer treatment, we applied the levels of nitrogen (N), potassium (K) and phosphorus (P) recommended for conventionally grown strawberry plants, i.e., 170, 140, and 100 kg/ha, respectively ^33,34^. The plants were fertilized three times per week with N and Ca (0.6 g/pot as calcium nitrate with total amount of 16.88 g/pot for 8 weeks), K and P (0.2 g/pot as di-potassium hydrogen phosphate with a total amount of 3.6 g/pot for 6 weeks) as well as Fe (0.1 g /pot as iron chelate EDDHA with a total amount of 2.4 g/pot for 8 weeks) dissolved in 100 ml of distilled water ^28^.

### Rearing of the herbivores *T. urticae* and *A. gossypii*

*Tetranychus urticae* (green form) used in experiments derived from a population reared on strawberry plants, *Fragaria x ananassa*. This spider mite population had been founded by specimens collected on whole common bean plants *Phaseolus vulgaris* L. ^28^ and was reared in the greenhouse at 26 ± 1°C, 65 ± 5% RH and natural daylight. For rearing, non-inoculated and unfertilized strawberry plants in the 3- to 4-leaf stage were periodically infested with mixed mobile stages of *T. urticae*, taken from other spider mite-infested strawberry plants. *Aphis gossypii* used in experiments was reared in a similar way on separate strawberry plants, *F.* x *ananassa,* but, different from the spider mites, the aphids had been originally collected from strawberry. Each plant/pot harboring one or the other herbivore was covered by a mesh cage (70 cm high, 30 cm Ø) to avoid contamination by other mites and insects. Both the spider mite and the aphid population were reared on strawberry for at least four generations (45 to 50 days) before use in the experiment.

### Experimental design

The aim of this experiment was to investigate changes in the level of defense by PGPR-inoculated plants (Ac, Ab and Pb treatments) in comparison to non-inoculated plants (chemical fertilization and control) against a later arriving herbivore (second release), when they had been previously infested by another herbivore (first release). To this end, strawberry plants (6 w old, in the 6- to 8-leaf stage) were randomly assigned to one of five rhizosphere treatments. Each pot with one strawberry plant constituted one experimental unit (replication). The experiment was a completely randomized 5×2 full factorial design with five rhizosphere treatments, i.e. Ab, Ac, Pb, chemical fertilization and control (n = 8 plants per treatment) crossed by two types of herbivore release (herbivore sequence), i.e., first infestation by aphids and second mites, and first infestation by mites and second aphids. In each rhizosphere treatment, 4 plants each were assigned to first release of mites or aphids. After each herbivore release, the population dynamics of each herbivore was monitored and the level of induced resistance of the infested plants was quantified by measuring the total phenol content.

### Population dynamics of *T. urticae* and *A. gossypii*

For mite release, 5 adult females were transferred with the aid of a moistened camel hair brush from the strawberry-reared stock population (reared on non-inoculated fertilized plants) to the abaxial surface of the youngest fully developed leaf of each potted strawberry plant. For aphid release, 4 adult females were transferred as stated previously. To warrant that the mites and aphids had successfully settled, we visually checked their fate after release. Infested leaves were marked to be specific for daily monitoring. To prevent movement of the herbivores among pots, each pot was covered by an acrylic cylinder (70 cm high × 30 cm diameter) closed on top by an organdy-mesh and kept under the afore-mentioned environmental conditions of the greenhouse. Four days after releasing the first herbivores, the youngest (5- to 6-d-old) fully developed non-infested leaf was clipped from each plant (n = 8 replicates for each treatment) at 8 am and placed immediately into a 15-ml centrifuge tube. The tubes were wrapped in aluminum foil and placed into a container filled with dry ice. The samples were weighed to the nearest mg using a digital balance (Sartorius GD503, Germany) and then frozen at −70°C until chemical analysis of the total phenol content.

Population development of *T. urticae* and *A. gossypii* on the potted strawberry plants was assessed once per week, with sampling starting 1 w after release of the first herbivore. To estimate mite abundance on each plant, the most densely populated leaf was detached, placed into a labeled polyethylene bag, transferred to the laboratory and there all mite life stages (including eggs) were counted using a stereo-microscope. The detached leaves were subsequently returned to the experimental unit they came from to allow the mites to re-integrate on their host plant. On each weekly sampling date, aphid abundance was determined on whole plants using a magnifying glass; on each date, the counting of the aphids was repeated three times and the mean of the three counts was used as final measure for this date. On the first sampling date (2022 June 9), that is, one week after releasing the first herbivore, the second herbivore release took place. To this end, 5 adult aphids were transferred from the rearing to the mite-infested plants and 5 adult mite females were transferred to the aphid-infested plants as described before. The abundances of all life stages of aphids and mites were weekly assessed over seven weeks, with sampling finishing on August 16, 2022. To take the second measure of the total phenol contents of leaves, 6 d after the second herbivore release, one fully-developed (5- to 6-d-old) non-infested leaf was clipped from each plant (n = 8 replicates for every treatment) and deep-frozen as described before.

### Measurement of total phenol contents of leaves

To determine the phenol concentration, 250 mg fresh leaves were crushed in liquid N2 with a pestle and mortar. After addition of 2.5 ml of 80% methanol, the extract was centrifuged at 4000 × g for 5 min. Then, 500 μl folin and 400 μl 20% sodium carbonate were added to 100 μl of the leaf extract. One ml of this solution was added to 5 ml of distilled water in test tubes, which were placed in a spectrophotometer (Labtron, LVS-A20) for 60 min at 765 nm. Total phenol was expressed as mg per g of dry leaf weight. A standard curve was obtained using different concentrations (0, 0.1, 0.2, 0.3, 0.4, 0.5, 1, 1.5) of galic acid (Acros Organic) ^35^.

### Measurement of total N and C contents of plants

To investigate the effect of the PGPR-treatments and herbivore-infestation on the level of the total N and C contents of the leaves after completion of the herbivore samplings, three newly expanded leaves (for both elements) were clipped from each of six plants from each treatment. The leaf samples were weighed to the nearest mg on a digital balance (Sartorius GD503, Germany). To measure the total N and C contents of leaves, the leaf samples were oven-dried at 70 °C for 48 h, and then weighed again on the digital balance. For each element (N and C), 6 replications (the same leaves for both N and C) per treatment were used. Each sample of dried plant material was then ground with a mortar and pestle and packed into tin capsules, each sample consisting of 2 to 3 mg. The N content was determined by the Kjeldahl method ^36^. The organic carbon was determined by Walkley-Black chromic acid wet oxidation method ^37^.

### Strawberry growth and physiological parameters

To characterize the effects of the PGPRs and herbivores on strawberry growth and physiology, the number of leaves and flowers and the photosynthesis rate and stomatal conductivity of the potted strawberry plants (6 w old, in the 6- to 8-leaf stage) were measured after completing the herbivore samplings. The leaf chlorophyll content and stomatal conductivity were measured by a SPAD portable leaf chlorophyll meter (SPAD-502, Minolta Camera, Co., Japan) and a leaf porometer system (model Sc-1), respectively. For both measurements, two newly expanded sunlit leaflets were randomly selected from each sampled plant. Each leaflet was one replication and measured once, and there were 12 replications for each rhizosphere treatment. After that, three leaves (above mentioned) were detached and their leaf area was estimated by Leaf Image Analysis System (WinDIAS, Delta-t company).

### Statistical analysis

Generalized estimating equations (GEE, autoregressive autocorrelation structure between sampling dates) were used to analyze the effect of the sequential release of the two herbivores (first aphids or first mites), the five rhizosphere treatments (Ab, Ac, Pb, chemical fertilization and control) and their interaction on population abundance of each herbivore. Data were log-transformed prior to analysis.

To assess the effect of rhizosphere treatment and herbivore sequence on growth and physiological parameters of strawberry plants, the data were analyzed by two-way ANOVAs. Significant differences between treatment pairs were determined by post hoc Tukey’s tests (P≤0.05). Before analysis, the data were checked for normality and homogeneity of variance using Kolmogorov-Smirnov and Bartlett tests, respectively. All data analyses were conducted using IBM SPSS 21 ^38^.

## Results

### Population dynamics of *T. urticae* and *A. gossypii*

Rhizosphere treatment, sequence of attack by mites and aphids and their interaction had significant effects on the abundance of each herbivore (Table 1). Irrespective of the rhizosphere treatment, aphids reared on mite-infested plants reached lower abundances than those first-released on clean plants (Figure 1). In chemically fertilized plants first-infested by aphids, the mean peak population reached 200 aphids per plant in the third week after release (Figure 1). On rhizobacteria-inoculated plants, mean aphid abundance was always lower than 40 individuals per plant in both herbivore sequences. The spider mites reached higher abundances when they were the first plant colonizers, regardless of rhizosphere treatment (180 and 100 individuals per leaf on chemically fertilized and control plants; Figure 2).

**Figure 1.**
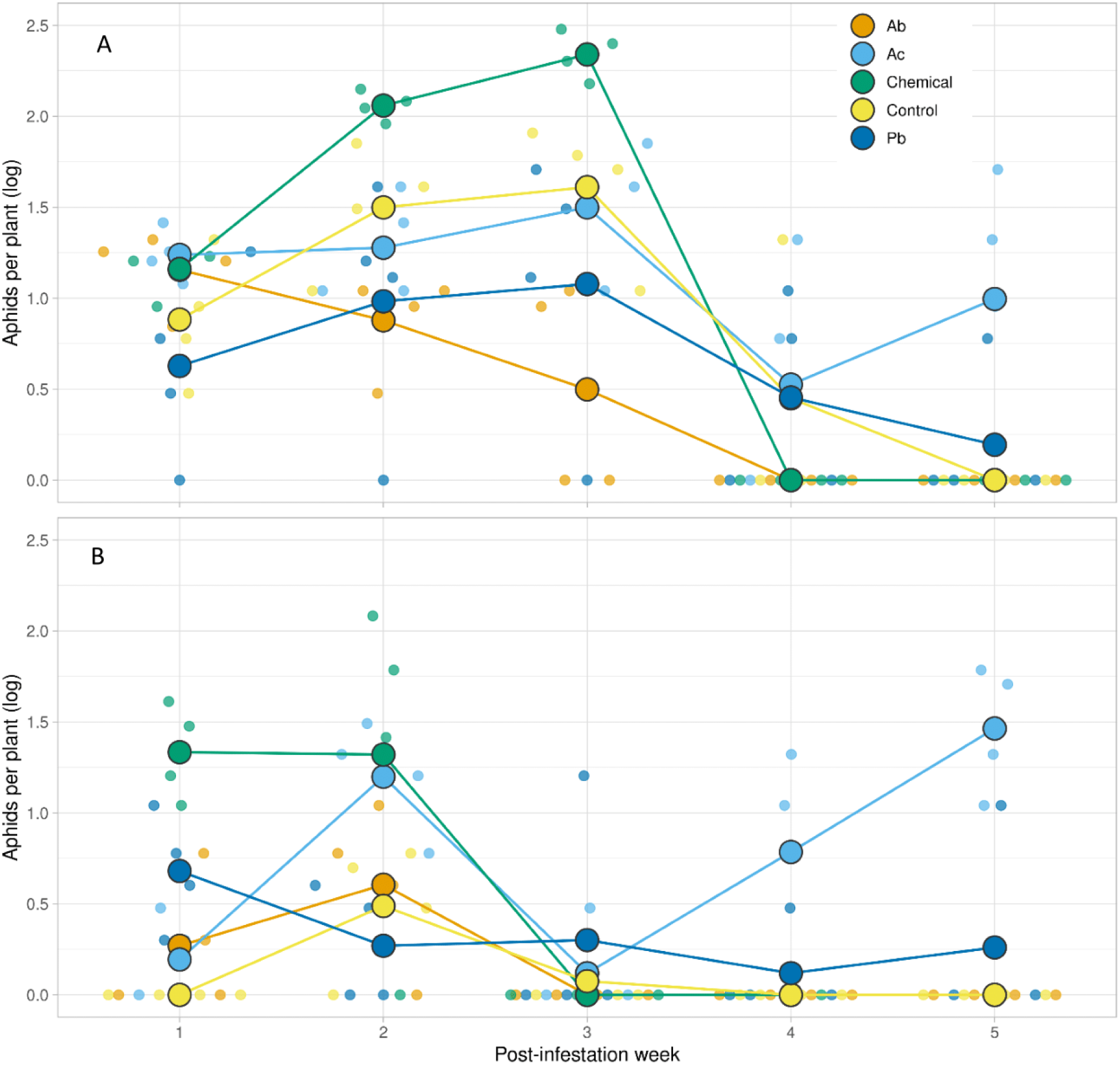
Mean abundance of cotton aphids *A. gossypii* (all developmental stages and adults) per strawberry plant in five rhizosphere treatments (Ab, Ac, Pb, chemical and control) in two herbivore sequences (A: aphids first, spider mites second; B: spider mites first, aphids second), during five sampling dates. Pale dots are the replicate data. Rhizosphere treatments: *Azotobacter chroococcum* (Ac), *Pseudomonas brassicacearum* (Pb), *Azospirillum brasilense* (Ab), no rhizosphere treatment (control), chemically fertilized (chemical). Statistical results are in table 1.

**Figure 2.**
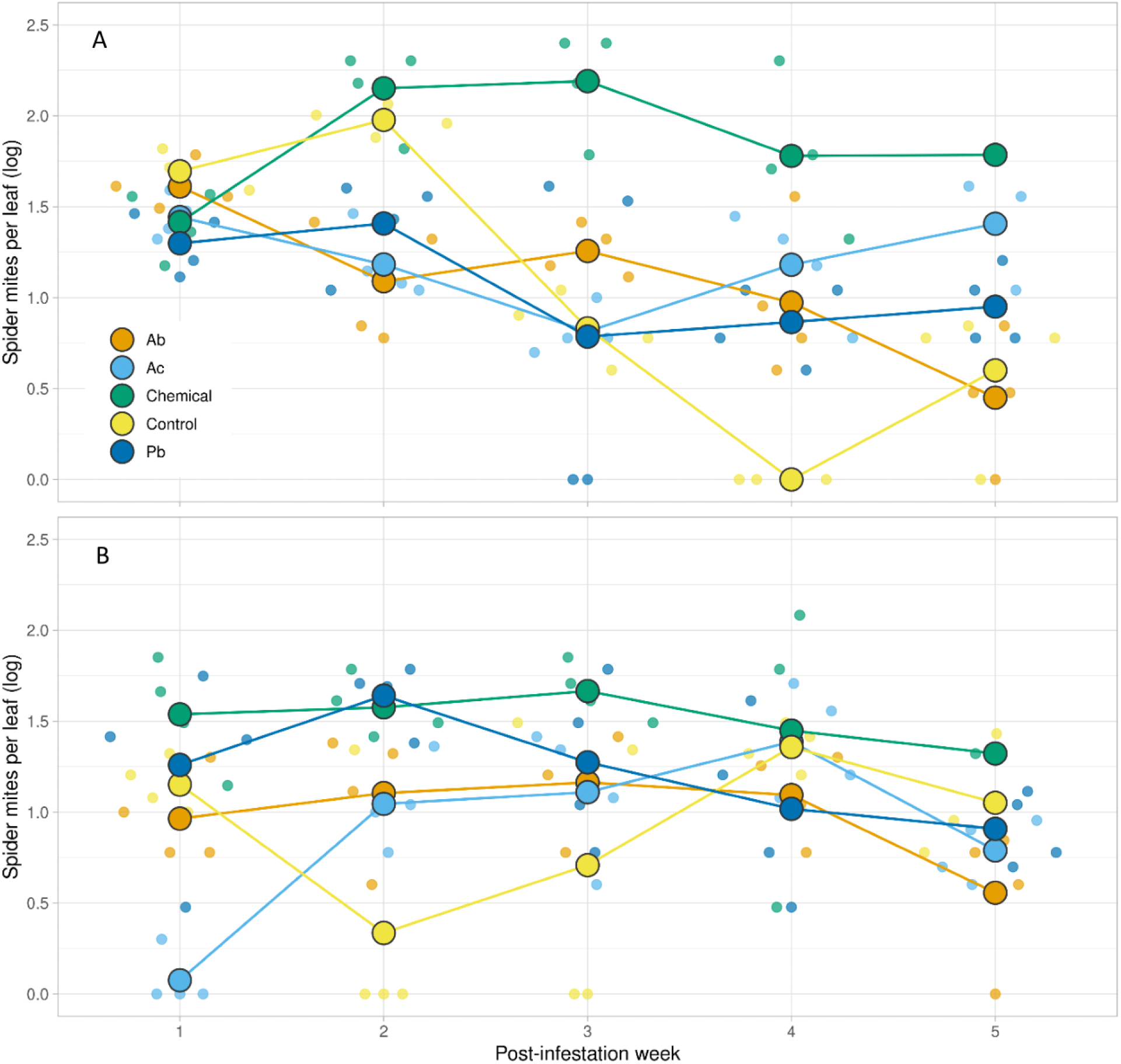
Mean abundance of two-spotted spider mites *T. uticae* (all developmental stages and adults) per strawberry leaf in five rhizosphere treatments (Ab, Ac, Pb, chemical and control) in two herbivore sequences (A: aphids first, spider mites second; B: spider mites first, aphids second), during five sampling dates. Pale dots are the replicate data. Rhizosphere treatments: *Azotobacter chroococcum* (Ac), *Pseudomonas brassicacearum* (Pb), *Azospirillum brasilense* (Ab), no rhizosphere treatment (control), chemically fertilized (chemical). Statistical results are in table 1.

**Table 1.** Results of separate GEEs on the effect of rhizosphere treatment (Ab, Ac, Pb, chemical fertilization and control), sequential attack by two herbivore species (herbivore sequence) and their interactions on population abundance of cotton aphids *A. gossypii* and two-spotted spider mites *T. urticae*.

| Source | Wald Chi-Square | df | P value |
| --- | --- | --- | --- |
| **Aphids** |  |  |  |
| Rhizosphere treatment | 86.337 | 4 | <0.001 |
| Herbivore sequence | 39.549 | 1 | <0.001 |
| Treatment * sequence | 16.370 | 4 | 0.003 |
| **Spider mites** |  |  |  |
| Rhizosphere treatment | 240.720 | 4 | <0.001 |
| Herbivore sequence | 16.817 | 1 | <0.001 |
| Treatment * sequence | 19.094 | 4 | 0.001 |

### Total phenol contents

After first herbivore release, leaf phenol content was significantly influenced by the interaction between rhizosphere treatment and herbivore sequence (Table 2, Figure 3). Total phenol content of plants infested by the first herbivore, before infestation by the second herbivore, significantly increased in rhizobacteria-inoculated plants in comparison to chemically fertilized and control plants. Total phenol contents were markedly higher in Pb-inoculated plants first-infested by spider mites than in the other treatments (Tukey; P≤0.5). In chemically fertilized plants, leaf phenol content was the lowest in both herbivore sequences. After the second herbivore release, leaf phenol content did not differ among rhizobacteria-inoculated and control plants and was the lowest in chemically fertilized plants (Tukey; P≤0.5; Figure 3).

**Figure 3.**
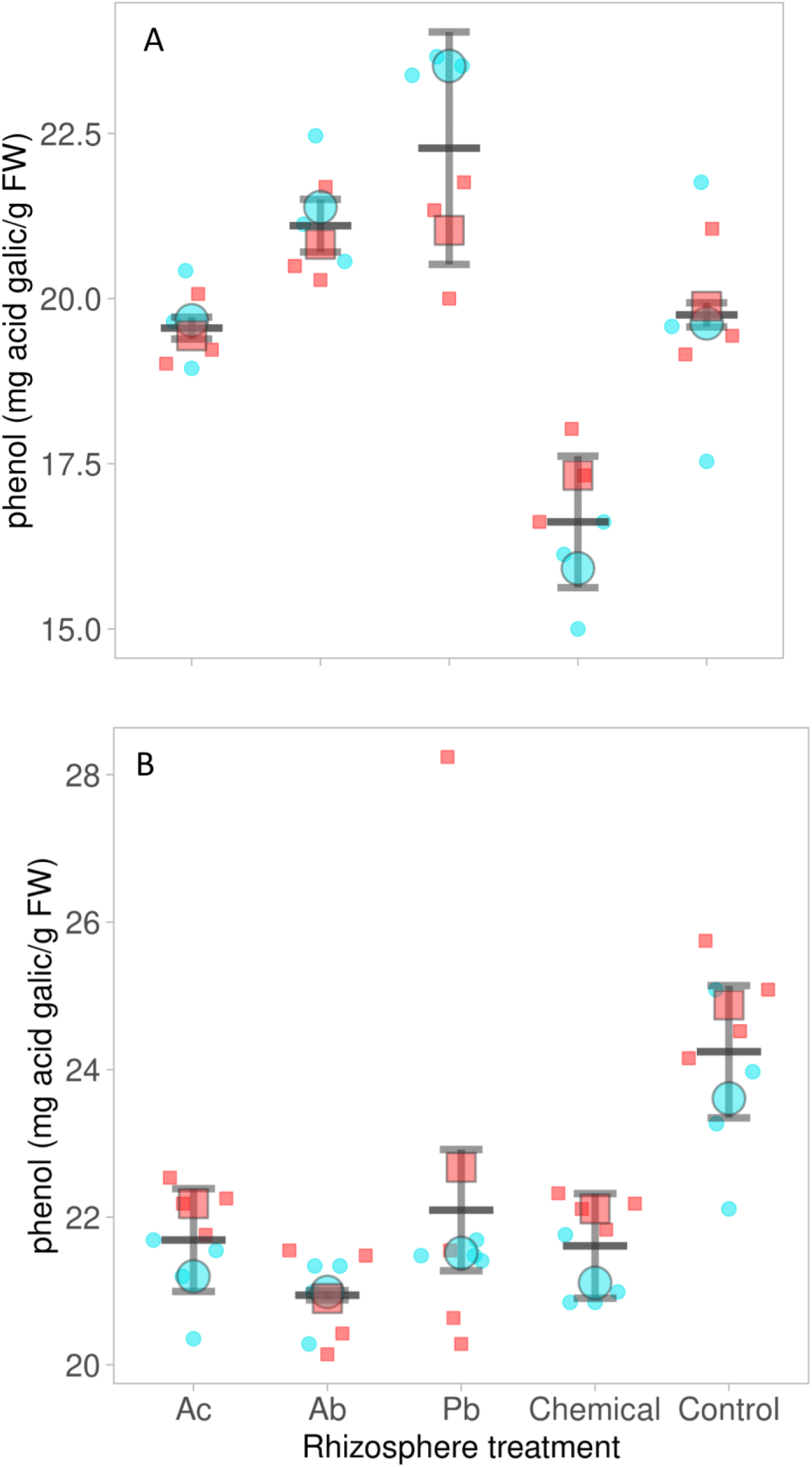
Mean (±SE) total phenol content of strawberry leaves in five rhizosphere treatments, 4 d after first release (A) and 6 d after second release (B), in two herbivore sequences (turquoise symbols for plants first-infested by aphids, red symbols for plants first-infested by mites). Pale dots are the replicate data. Rhizosphere treatments: *Azotobacter chroococcum* (Ac), *Pseudomonas brassicacearum* (Pb), *Azospirillum brasilense* (Ab), no rhizosphere treatment (control), chemically fertilized (chemical). Statistical results are in table 2.

**Table 2.** Results of separate ANOVAs on the effects of rhizosphere treatment (Ab, Ac, Pb, chemical fertilization and control), sequential attack by two herbivore species (herbivore sequence) and their interactions on leaf total phenol content after first and second herbivore release.

| Source | df | F | P value |
| --- | --- | --- | --- |
| **After first release** |  |  |  |
| Rhizosphere treatment | 4 | 14.124 | <0.001 |
| Herbivore sequence | 1 | .697 | 0.414 |
| Treatment * sequence | 4 | 9.839 | <0.001 |
| **After second release** |  |  |  |
| Rhizosphere treatment | 4 | 3.591 | 0.017 |
| Herbivore sequence | 1 | 2.125 | 0.155 |
| Treatment * sequence | 4 | .169 | 0.953 |

### Strawberry growth, physiological and chemical parameters

The photosynthesis rate was significantly influenced by rhizosphere treatment and was higher in Ac- and Pb-inoculated plants than in control and chemically fertilized plants (Table 3, Tukey: P≤0.5; Figure 4). Stomatal conductivity (gs) of strawberry plants was influenced by rhizosphere treatment and herbivore sequence and was the highest in Ac- and Pb-inoculated plants. Also, in plants first-infested by aphids, stomatal conductivity incrased significantly in comparison to those first-infested by mites (t=-2.32 and P<0.05, Figure 4). The number of leaves and flowers, leaf area, N and C were siginificantly influenced by the interaction between rhizosphere treatment and herbivore sequence (Table 3). In both herbivore sequences, Ac- and Pb-inoculated plants had more leaves than plants in the other rhizosphere treatment/herbivore sequence combinations (Figure 5). However, in Ac-inoculated, chemically fertilized and control plants first attacked by aphids, the number of leaves was significantly higher than in the corresponding plants first-infested by spider mites (Figure 5). Leaf area significantly decreased in control plants in both herbivore sequences compared to the other rhizosphere treatment/herbivore sequence combinations (Tukey: P≤0.5; Figure 5). In both herbivore sequences, Pb-inoculated plants produced the most flowers. In Ac-treated and control plants first-infested by aphids, the number of flowers was higher than in the corresponding plants first-infested by spider mites, in Ab-treated plants the reverse was the case. Chemically fertilized plants did not produce any flowers (Figure 5). Leaf nitrogen content reached the highest level in Pb-inoculated plants first attacked by aphids (Figure 6). All rhizosphere treatments except the control had higher leaf nitrogen contents when first attacked by aphids than when first attacked by spider mites. Ac-inoculated plants, in both herbivore sequences, had the highest C content (Tukey: P≤0.5; Figure 6).

**Figure 4.**
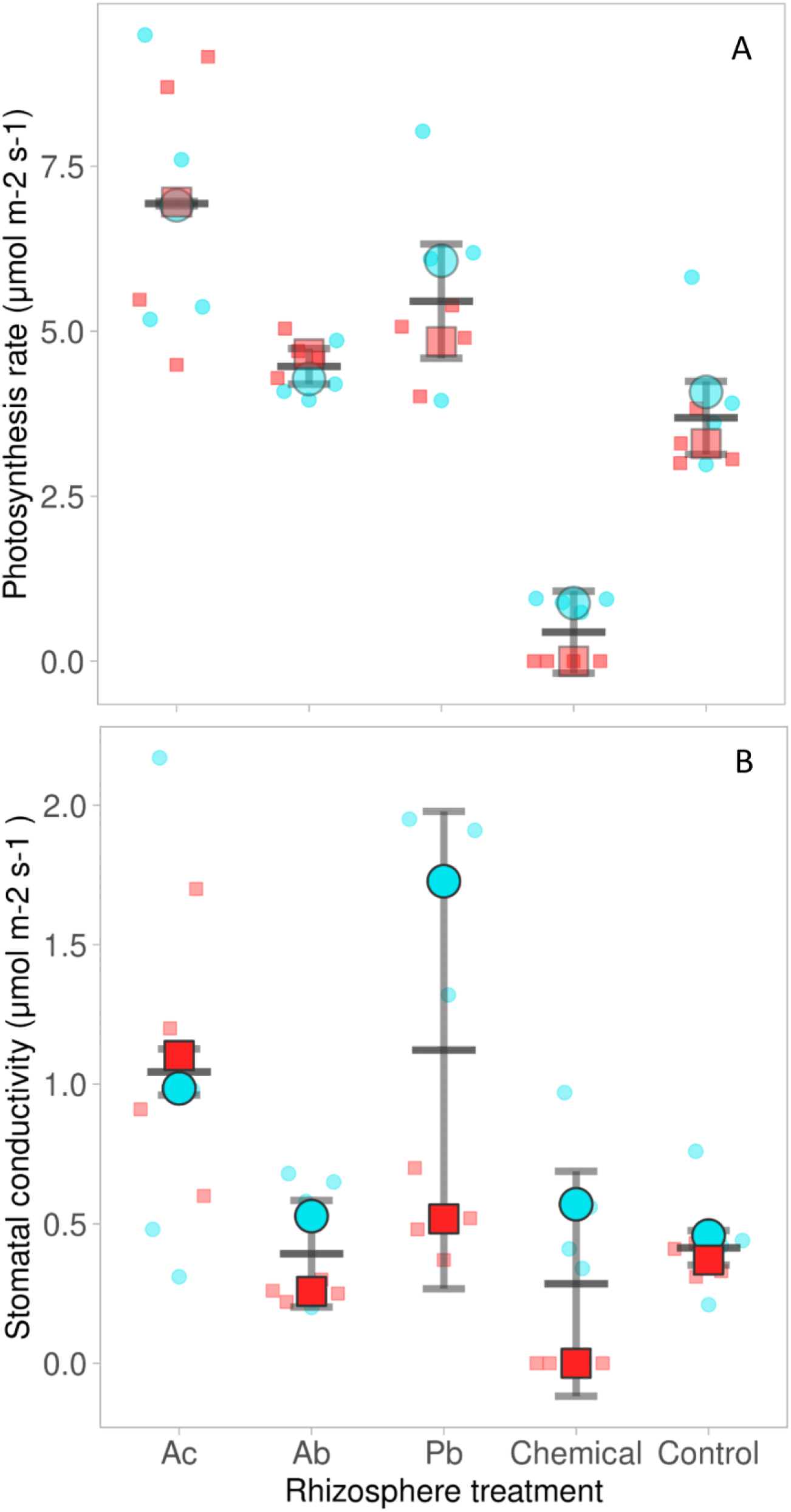
Mean (±SE) photosynthesis rate (A) and stomatal conductivity (B) of strawberry plants in five rhizosphere treatments and two herbivore sequences (turquoise symbols for plants first-infested by aphids, red symbols for plants first-infested by mites). Pale dots are the replicate data. Rhizosphere treatments: *Azotobacter chroococcum* (Ac), *Pseudomonas brassicacearum* (Pb), *Azospirillum brasilense* (Ab), no rhizosphere treatment (control), chemically fertilized (chemical). Statistical results are in table 3.

**Figure 5.**
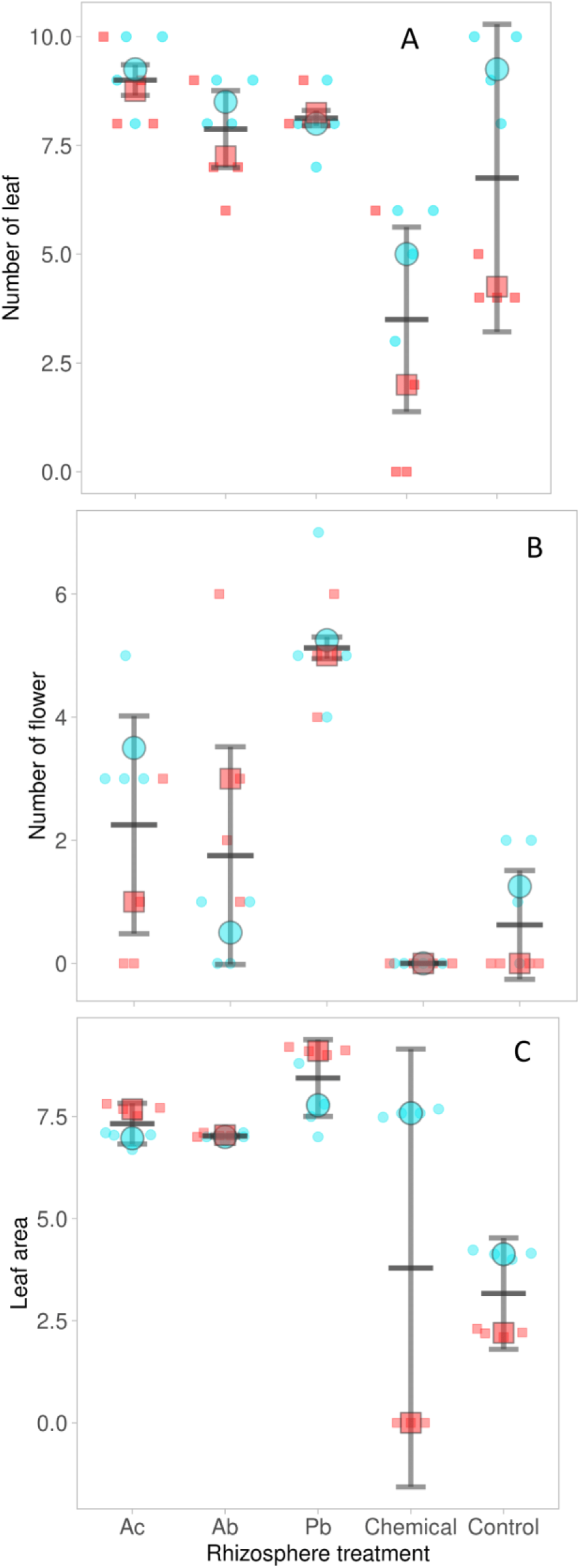
Mean (±SE) number of leaves (A) and flowers (B), and mean leaf area (C) in five rhizosphere treatments in two herbivore sequences (turquoise symbols for plants first-infested by aphids, red symbols for plants first-infested by mites). Pale dots are the replicate data. Rhizosphere treatments: *Azotobacter chroococcum* (Ac), *Pseudomonas brassicacearum* (Pb), *Azospirillum brasilense* (Ab), no rhizosphere treatment (control), chemically fertilized (chemical). Statistical results are in table 3.

**Figure 6.**
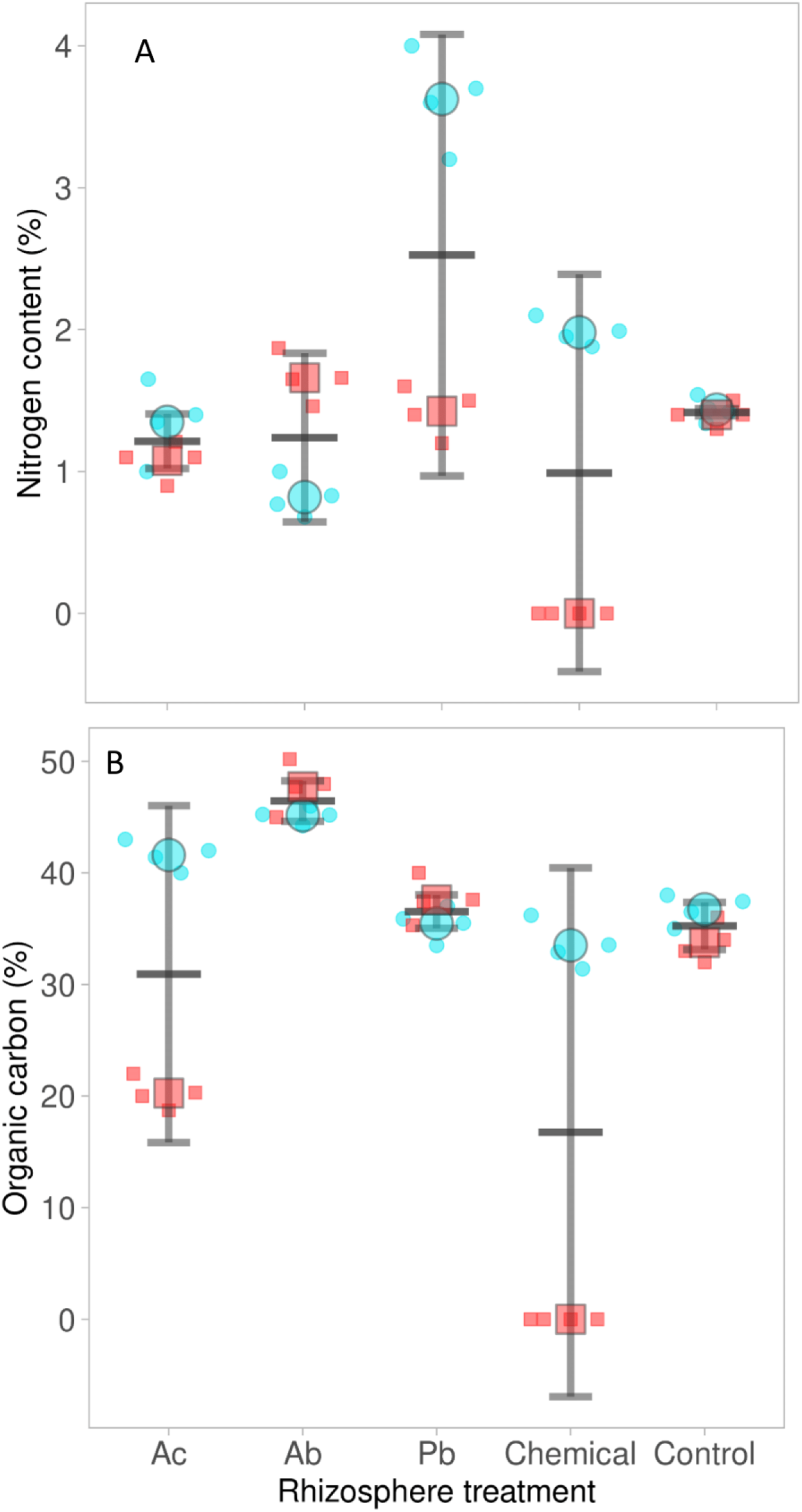
Mean (±SE) percent nitrogen (A) and organic carbon (B) of strawberry leaves in five rhizosphere treatments in two herbivore sequences (turquoise symbols for plants first-infested by aphids, red symbols for plants first-infested by mites). Pale dots are the replicate data. Rhizosphere treatments: *Azotobacter chroococcum* (Ac), *Pseudomonas brassicacearum* (Pb), *Azospirillum brasilense* (Ab), no rhizosphere treatment (control), chemically fertilized (chemical). Statistical results are in table 3.

**Table 3.** Results of separate ANOVAs on the effects of rhizosphere treatment (Ab, Ac, Pb, chemical fertilization and control), sequential attack by two herbivore species (herbivore sequence) and their interactions on photosynthesis, stomatal conductivity, number of leaves and flowers, leaf area, nitrogen content and organic carbon.

| Source | df | F | Sig. |
| --- | --- | --- | --- |
| **Photosynthesis** |  |  |  |
| Rhizosphere treatment | 4 | 15.36 | <0.001 |
| Herbivore sequence | 1 | 1.29 | 0.27 |
| Treatment * sequence | 4 | 0.27 | 0.89 |
| **Stomatal conductivity** |  |  |  |
| Rhizosphere treatment | 4 | 5.07 | 0.006 |
| Herbivore sequence | 1 | 5.56 | 0.019 |
| Treatment * sequence | 4 | 2.13 | 0.115 |
| **Number of leaves** |  |  |  |
| Rhizosphere treatment | 4 | 12.49 | <0.001 |
| Herbivore sequence | 1 | 12.55 | 0.002 |
| Treatment * sequence | 4 | 2.96 | 0.045 |
| **Number of flowers** |  |  |  |
| Rhizosphere treatment | 4 | 10.76 | <0.001 |
| Herbivore sequence | 1 | 0.37 | 0.548 |
| Treatment * sequence | 4 | 3.61 | 0.056 |
| **Leaf area** |  |  |  |
| Rhizosphere treatment | 4 | 32.6 | 0.001 |
| Herbivore sequence | 1 | 16.56 | 0.002 |
| Treatment * sequence | 4 | 21.1 | 0.001 |
| **Nitrogen content** |  |  |  |
| Rhizosphere treatment | 4 | 14.7 | 0.001 |
| Herbivore sequence | 1 | 7.30 | 0.018 |
| Treatment * sequence | 4 | 5.87 | 0.015 |
| **Organic carbon** |  |  |  |
| Rhizosphere treatment | 4 | 20.81 | <0.001 |
| Herbivore sequence | 1 | 4.21 | 0.020 |
| Treatment * sequence | 4 | 3.88 | 0.020 |

## Discussion

Herbivores with different feeding modes attacking the same plant can exert strong direct and indirect effects on each other ^3,39^ and above-ground herbivore interactions may be mediated by plant-associated rhizosphere micro-organisms. Our study documents that sequential attack of strawberry plants by spider mites and cotton aphids, regardless of the herbivore sequence, adversely affects the abundance of the later arriving herbivore. The herbivore-interactions were exclusively mediated by the host plant’s systemic defense, because the spider mites and aphids feed on different parts of the plants and exploit different plant tissues and resources (parenchyma cell contents and phloem sap, respectively). Similarly, Kiełkiewicz et al. ^40^ showed that, regardless of which attacker was first, priming of *Arabidopsis* plants by spider mites or aphids decreased the reproductive performance of the later arriving herbivore species. Strikingly, in our study, the herbivores’ abundances were much higher and the mutual adverse effects were much stronger in chemically fertilized plants than in plants associated with PGPRs.

Two-spotted spider mites have a greater population growth capacity and plant colonization ability than cotton aphids ^39^. Accordingly, our study revealed that spider mite presence on the plants had a much stronger suppressing effect on the aphid population than aphid presence had on mite population dynamics. Mite abundance on Pb-inoculated plants that were initially infested by aphids, which had the highest leaf nitrogen content among rhizosphere treatments, reached 50 individuals per strawberry leaf on the third sampling date, which was the highest abundance on this date among all rhizosphere treatments and herbivore attack sequences. Alizade et al. (2016) showed that the population growth rate of two-spotted spider mites increases with increasing leaf N content of strawberry. In contrast, initial mite infestation strongly suppressed aphid population growth (fewer than 25 individuals per plant) in all rhizosphere treatments.

Rhizobacteria can enhance plant resistance to aboveground-living herbivorous insects and mites by inducing systemic resistance (ISR) ^42,43^. ISR is considered a plant priming mechanism, which enhances the biosynthesis of defense-related chemical compounds such as phenols, flavonoids and other secondary metabolites regulated by the JA and/or SA pathways ^4,27,44^ For example, colonization of rice roots by *Pseudomonas fluorescens* may induce ISR, which enhances the accumulation of phenolic compounds in green plant parts ^45^. Our study shows that rhizobacteria inoculation primed the strawberry plants’ defense mechanisms to attack by spider mites or aphids, enabling the plants to produce higher amonts of secondary metabolites such as total phenols. As a consequence, the abundances of both herbivores were significantly lower on rhizobacteria-inoculated plants compared to chemically fertilized and control plants.

The feeding activity by two-spotted spider mites is well known for inducing defense mechanisms and upregulation of primarily JA, but also SA, pathways in various plant species including strawberry, which also causes the emission of volatile organic compounds (VOCs) to attract the herbivores’ natural enemies ^46–48^. Theses VOCs may be eavesdropped by other herbivores and may disrupt the attractiveness of spider mite-infested plants to aphids ^49^. Several studies showed that aphids produce and secrete effectors that modulate the plants’ defense response to herbivores and pathogens ^50,51^. For example, feeding by green peach aphids *Myzus persicae* was found to induce the expression of SA-signaling pathway marker genes such as PR-1 ^52^. The effects of induced defense by the herbivores were most pronounced in chemically fertilized plants, in which preempting mite or aphid populations strongly suppressed the heterospecific herbivore population. The populations of both *T. urticae* and *A. gossypii* as first colonizers exploded on chemically fertilized plants and reached more than 150 mites per leaf and 200 aphids per plant. Conversely, priming of the systemic defense system of strawberry plants by rhizobacteria inoculation prevented the mite and aphid population as second colonizers to reach plant-damaging abundances. The peak population of *T. urticae* on chemically fertilized and rhizobacteria-inoculated plants initially infested by aphids were approximately 60 and 25 mites per leaf, respectively, and a similar trend was found for the peak populations of *A. gossypii* per plant on strawberry plants first-infested by spider mites. These patterns suggest that the defense system of strawberry plants triggered by rhizobacteria has a stronger effect on the population dynamics of later arriving herbivores than the plant defense system elicited by initial presence of aphid or mite population in rhizobacteria absence. Future research should address the molecular basis of the differences between rhizobacteria- and herbivore-triggered plant defense mechanisms and their interplay.

In addition to activating the defense system of plants against future attacks by herbivores and/or pathogens, rhizobacteria also assist the plants in nutrient acquisition through a variety of mechanisms, including N-fixation, enhancing nutrient bioavailability through phosphate-solubilization, producing siderophores for Fe chelation and increasing solubilization of other micronutrients ^53,54^. Enhanced stomatal conductivity and photosynthesis rate along with a larger leaf area and higher leaf number in Pb- and Ac-inoculated plants suggests that Pb and Ac increased the plants’ efficiency in transport and distribution of water and nutrients such as soluble proteins, assimilates, and hormones ^55^. High infestation levels of spider mites and aphids on chemically fertilized plants greatly decreased stomatal conductivity. Moreover, Pb and Ac enhanced flower production and probably future fruit production compared to the other rhizosphere treatments. It seems that Pb and Ac inoculation, by inducing defense mechanisms of the strawberry plants including elevating the total phenol contents, can prevent explosive growth of the populations of *T. urticae* and *A. gossypii*, irrespective of the herbivore attack sequence. Similarly, separate inoculation of strawberry plants with two bacterial isolates, *Bacillus amylolequefaciens* BChi1 and *Paraburkholderia fungorum* BRRh-4, significantly increased vegetative growth, yield, contents of various antioxidants and total antioxidant activities such as total phenolic content ^56^. Irrespective of the rhizosphere treatment, stomatal conductivity of plants first infested by spider mites was significantly lower than that of plants first infested by aphids. Also, first infestation by spider mites had, on average, stronger negative effects on other plant physiological parameters than first infestation by aphids. For example, chemically fertilized plants first infested by spider mites were completely defoliated by the fifth sampling date. Spider mites typically pierce the lower surface of strawberry leaves to suck fluids from mesophyll cells and then secondary damage occurs to epidermal cells because of mesophyll dehydration. Epidermal cells, including stomatal guard cells, consequently dehydrate, which results in closing the stomatal pores and preventing gas exchange. Moreover, tissue dehydration caused by spider mite feeding can reduce the concentration of soluble proteins ^55,57^. These results agree with those of Sances et al. ^58^, who showed a negative correlation between *T. urticae* density and stomatal conductivity on strawberry plants.

In conclusion, we demonstrated that spider mites and cotton aphids have mutual adverse effects on each other, especially on chemically fertilized plants. However, priming of plant defense mechanisms by rhizobacteria inoculation through increased production of total phenol contents, significantly decreased population growth of later arriving aphids or spider mites, regardless of the herbivore attack sequence, compared to chemically fertilized and control plants. Regarding the herbivore sequence, initial infestation by spider mites had stronger adverse effects on the physiologicy of the strawberry plants than initial infestation by aphids. For example, the N content of Pb-inoculated strawberry plants was significantly higher when first-infested by *A. gossypii* than when first-infested by *T. urticae*. Our study also suggests that inoculation of strawberry plants by rhizobacteria (ISR) induced stronger adverse plant-mediated effects on aboveground herbivore populations than the plant defense mechanisms activated by attack by aphid or mite populations (IR) in rhizobacteria absence. Overall, our study emphasizes that rhizobacteria inoculation not only mitigates the damage caused by sequentially attacking herbivores, through increased total phenol content of the plant, but also amends the nutritional status and photosynthesis activity of strawberry plants. It may well be that rhizobacteria inoculation also enhances the recruitment of third trophic level natural enemies by the plants, through changing the emission of herbivore-induced plant volatiles (HIPVs) and/or the nutritional quality of the herbivores as prey or hosts ^12,13,31,48^. This issue remains to be addressed in future studies. In order to promote the production of healthy fruits free of pesticide residues, future research should scrutinize the consequence of rhizobacteria application in integration with other sustainable plant protection methods such as the use of natural enemies.

## Author contributions

AH and MH conceived the study idea, AH, MH and PS designed the experiments. AH conducted the experiments, analyzed the data, and wrote the first draft of the manuscript. MH contributed to methods development and managed the project locally. PS contributed to results analysis and presentation. MH and PS contributed to writing. All authors contributed to manuscript revision, read, and approved the submitted version.

## Data availability

The datasets generated during the current study are available from the corresponding authors on reasonable request.

## Consent for publication

All authors consent to the publication of this manuscript in the Journal of Pest Science.

## Acknowledgments

The research deputy of the Ferdowsi University of Mashhad is acknowledged for granting the use of their facilities.

